# Modeling human age-associated increase in Gadd45γ expression leads to spatial recognition memory impairments in young adult mice

**DOI:** 10.1101/2020.01.12.903112

**Authors:** David V.C. Brito, Kubra Gulmez Karaca, Janina Kupke, Franziska Mudlaff, Benjamin Zeuch, Rui Gomes, Luísa V. Lopes, Ana M.M. Oliveira

## Abstract

Aging is associated with the progressive decay of cognitive function. Hippocampus-dependent processes, such as the formation of spatial memory, are particularly vulnerable to aging. Currently, the molecular mechanisms responsible for age-dependent cognitive decline are largely unknown. Here, we investigated the expression and function of the growth arrest DNA damage gamma (Gadd45γ) during aging and cognition. We report that Gadd45γ expression is increased in the hippocampus of aged humans and that Gadd45γ overexpression in the young adult mouse hippocampus compromises cognition. Moreover, Gadd45γ overexpression in hippocampal neurons disrupted CREB signaling and the expression of well-established activity-regulated genes. This work shows that Gadd45γ expression is tightly controlled in the hippocampus and its disruption may be a mechanism contributing to age-related cognitive impairments observed in humans.

## 1. Introduction

Age-related cognitive decline in humans affects about 40% of individuals aged 65 years or older (Aigbogun et al., 2017), even though deterioration of cognitive functions may start earlier (Singh-Manoux et al., 2012). Long-term memory formation requires activity-regulated signaling that results in *de novo* gene expression. These genomic responses are known to be disrupted in the aged hippocampus (Stefanelli et al., 2018). Therefore, age-associated changes that dysregulate the coupling between neuronal activity and gene transcription likely underlie age-related cognitive deficits.

Recently, we and others showed that the growth arrest DNA damage gamma (Gadd45γ) is required for memory formation in the prelimbic prefrontal cortex (Li et al., 2018) and the hippocampus (Brito et al., 2020). Moreover, we found that aging reduces Gadd45γ expression in the mouse hippocampus and that mimicking this reduction in young adult mice induces age-like memory impairments (Brito et al., 2020). At the molecular level, Gadd45γ is required for cAMP response element-binding protein (CREB) activation in response to neuronal activity and associated gene expression (Brito et al., 2020). In the current study, we found that in *postmortem* human hippocampal tissue from aged individuals, Gadd45γ expression levels are increased relative to young donors. Furthermore, we showed that increasing Gadd45γ levels in the mouse hippocampus led to impairments in memory formation and disrupted CREB activation and memory-related gene expression in cultured hippocampal neurons.

Overall, this work together with our previous findings (Brito et al., 2020), demonstrate a requirement for tight regulation of neuronal Gadd45γ levels in gene expression regulation and cognitive abilities. Thus, dysregulation of Gadd45γ expression might be an underlying mechanism involved in age-related cognitive impairments observed in mice and humans.

## 2. Materials and Methods

### 2.1. Subjects

The use of human samples was conducted in accordance with the Helsinki Declaration as well as national ethical guidelines. Protocols were approved by the Local Ethics Committee and the National Data Protection Committee. The biospecimens were obtained 36h *postmortem* from healthy aged (60–65 years old) and young (21–22 years old) individuals (Temido-Ferreira et al., 2018). The tissue was processed and preserved for molecular analyses as previously described (Pliassova et al., 2016). Young adult male C57BL/6N mice (Charles River, Sulzfeld, Germany) were 3-months-old at the time of behavior experiments. Mice were group-housed (3-4 mice per cage) on a 12h light/dark cycle (22 ± 1°C, 55 ± 10% relative humidity) with *ad libitum* access to water and food. All behavioral experiments took place during the light phase. All procedures were carried out in accordance with German guidelines for the care and use of laboratory animals and with the European Community Council Directive 86/609/EEC.

### 2.2. Recombinant adeno-associated virus (rAAVs)

Viral particles were produced and purified as described previously (Gulmez Karaca et al., 2020). Overexpression of Gadd45γ was achieved by using a viral vector that contained the mouse CamKIIα promoter upstream of the Gadd45γ full-length mouse cDNA sequence. As a control vector, we used a construct containing the CamKIIα promoter driving the expression of GFP. For each virus batch, toxicity was analyzed on primary hippocampal cultures before the start of the experiments. For this, different regions of the coverslip were imaged using identical microscope settings and the number of dead cells was quantified using Fiji (Schindelin et al., 2012) on day *in vitro* (DIV) 10.

### 2.3. Primary hippocampal cultures

Hippocampal cultures from newborn C57Bl/6N mice (Charles River, Sulzfeld, Germany) were prepared and maintained as previously described (Gulmez Karaca et al., 2020). rAAV infection of cultures occurred on DIV 4. Experiments were performed on DIV 10. To induce action potential bursting, cultures were treated with 50 μM bicuculline (Enzo Life Sciences, Germany).

### 2.4. Stereotaxic surgery

rAAVs were injected into the dorsal hippocampus at the following coordinates relative to Bregma: − 2 mm anteroposterior, ± 1.5 mm medio-lateral, − 1.7, − 1.9 and − 2.1 mm dorsoventral. A total volume of 1.5 μl was injected per hemisphere at 200 nl/min. Following injections at each individual site, the needle was left in place for 60s. Behavioral experiments started 2 weeks after rAAVs delivery. After behavioral testing, histological analysis was performed to confirm tissue and cellular integrity.

### 2.5. Behavioral testing

Before behavioral testing started, mice were habituated to the experimenter and behavioral room by handling for 3 consecutive days, 1 minute per mouse. Object-location test and contextual fear conditioning were performed as previously described (Oliveira et al., 2012; Oliveira et al., 2016). The open field test was carried out within the first session of the object-place recognition training as previously described (Gulmez Karaca et al., 2018).

### 2.6. Quantitative reverse-transcription PCR

Total RNA from human tissue was extracted and cDNA produced as previously described (Temido-Ferreira et al., 2018). For RNA isolation from mouse hippocampal tissue, the tissue was rapidly dissected, placed in RNAlater (Sigma, Munich, Germany) and isolated using the RNeasy Plus Mini Kit (Qiagen, Hilden, Germany) with additional on-column DNase I digestion, according to the manufacturer’s instructions. cDNA production and quantitative reverse-transcription PCR (q-RT-PCR) was performed as previously described (Brito et al., 2020). The following TaqMan probes were used: *Arc* (Mm00479619_g1), *c-Fos* (Mm00487425_m1), *FosB* (Mm00500401_m1), *Gadd45*α (Mm00432802_m1), *Gadd45*β (Mm00345123_m1), *Gadd45*γ (Mm00442225_m1), *Egr1* (Mm00656724_m1) and *Npas4* (Mm00463644_m1). For human genes the following TaqMan probes were used: *Gadd45*α (Hs00169255_m1), *Gadd45*β (Hs00169587_m1), *Gadd45*γ (Hs00198672_m1). Expression levels of target genes were normalized to the expression of the housekeeping gene *GusB* (Mm00446953_m1) or β-actin (Hs01060665_g1) for mouse or human genes, respectively. Controls were used to exclude the possibility of DNA or RNA contaminations.

### 2.7. Western blotting

Western blotting was performed as previously described with minor modifications (Brito et al., 2020; Gulmez Karaca et al., 2020). Briefly, hippocampal cultures infected on DIV 4 were lysed on DIV 10 in SDS sample buffer. In the case of western blotting of tissue samples, the dorsal hippocampus was microdissected from mouse brain and homogenized in RIPA buffer (150 mM NaCl, 1% Triton X-100, 0.5% sodium deoxycholate, 0.1% SDS, 50 mM Tris, pH 8.0) supplemented with 1% protease inhibitor cocktail (Sigma-Aldrich, Munich, Germany) and 1% phosphatase inhibitor cocktail II and III (Sigma-Aldrich, P5726, P0044), the whole procedure was performed at 4°C. Protein concentration was measured by Bradford assay and 20 µg of protein was loaded on a 10% polyacrylamide gel after being denatured at 95 °C for 5 min. After SDS-PAGE, gels were blotted onto a nitrocellulose membrane (GE Healthcare, Buckinghamshire, UK) and later blocked in 5% milk and probed with the following antibodies: phospo-CREB (1:6000, Millipore #05-667), total-CREB (1:5000, Cell Signaling, #4820), β-Actin (1:1000, Santa Cruz, #SC-47778) or α-Tubulin (1:40000, Sigma-Aldrich, #T9026). Antibodies were diluted in 5% milk in PBS-T (total-CREB, α-Tubulin and β-Actin) or in 5% bovine serum albumin in PBS-T (phospho-CREB). Next, the membranes were incubated with horseradish peroxidase-conjugated secondary antibodies and later analyzed using a ChemiDoc™ Imaging System (Bio-Rad). Data is presented as ratio of phosphorylated/total protein normalized internally to each uninfected condition.

### 2.8. tatistical information

For normally distributed data sets, two-tailed unpaired Student’s t test or one-way ANOVA were used to compare two or more groups respectively (significant data is marked with *). Two-tailed Mann-Whitney test was used to compare two distinct groups for non-Gaussian distribution (significant data is marked with #). Correlation analysis was performed using Pearson correlation coefficient or Spearman correlation for normally distributed or non-parametric data, respectively. The sample size was determined based on similar experiments carried out in the past. All plotted data represent mean ± SEM. Statistics were performed using GraphPad Prism for Mac OS X, version 8. For behavioral experiments the investigators were blind to group allocation during data collection and analysis. For *in vitro* experiments no blinding was performed since the outcome was dependent on software analysis and not manual scoring.

## 3. Results

### 3.1. Aging increases Gadd45γ expression in the human hippocampus

Aberrant gene expression patterns are an evolutionarily conserved hallmark of aging. However, no overall correlation between age-associated gene expression in mice and humans has been detected (Zahn et al., 2007). We asked whether Gadd45γ expression in human aged hippocampus would be compromised as observed in mice (Brito et al., 2020). We analyzed the expression of Gadd45 family members in young and aged human hippocampi as we previously described (21–65 years old) (Temido-Ferreira et al., 2018) (Figure 1A). We did not find any correlation between age and *Gadd45*α expression. Interestingly, we found that hippocampal *Gadd45*β and *Gadd45*γ levels were increased (∼4.8 and ∼8.6 fold, respectively) as age progressed. This result, together with our previous findings in aged mouse tissue (Brito et al., 2020), suggests that age-related Gadd45 expression changes in the hippocampus may not be conserved in mice and humans.

**Figure 1.**
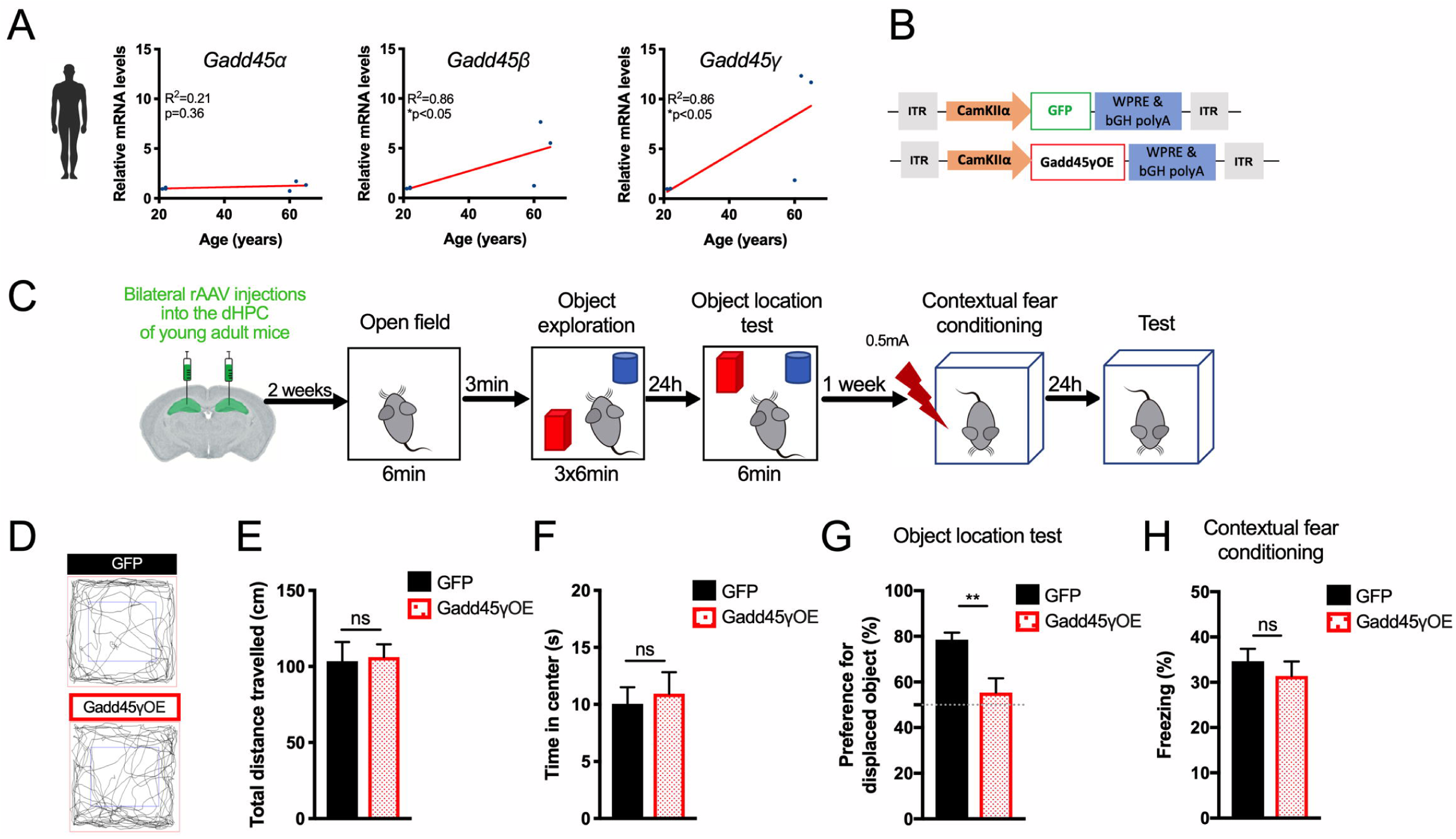
Human hippocampal Gadd45γ expression is increased during aging and overexpressing Gadd45γ in the mouse hippocampus impairs object location memory. **A)** Correlational analysis between the expression of *Gadd45*α, *Gadd45*β and Gadd45γ in human *postmortem* hippocampal tissue and donors’ age (N=6). Correlation analysis was performed using Pearson correlation coefficient or Spearman correlation. **B)** Schematic representation of the viral constructs used. The viral vector contains a CamKIIα promoter driving Gadd45γ overexpression (Gadd45γOE) or GFP as a control (GFP). **C)** Schematic representation of the experimental design for behavioral tests. **D)** Representative exploration patterns of all groups during open field test. **E)** Locomotion analysis of the different groups measured as the total distance travelled during the open field test (N=8-9). **F)** Anxiety-like behavior analysis measured as percentage of time spent in the center of the arena during the open field test (N=8-9). **G)** 24h object location memory test of young adult mice expressing GFP or Gadd45γOE in the dHPC (N=13-15). Dashed line represents equal preference for either object (chance preference). **H)** 24h contextual fear memory test of young adult mice expressing GFP or Gadd45γOE in the dHPC (N=9). ns: not significant; **p<0.01 by two-tailed unpaired Student’s t-test. Error bars represent SEM.

### 3.2. Gadd45γ overexpression leads to impairments in spatial recognition memory

Next, we sought to model the human aging-associated Gadd45γ increase in the mouse hippocampus and determine the cellular and behavioral consequences of neuronal Gadd45γ overexpression. Given that previous studies showed a selective function for Gadd45γ in memory formation (Brito et al., 2020; Li et al., 2018), we focused on Gadd45γ. We stereotaxically delivered a viral vector containing the mouse CamKIIα promoter driving the expression of Gadd45γ, or GFP as a control, into the dorsal hippocampus (dHPC) of young adult mice (Figure 1B,C). We validated viral expression in the dHPC of injected animals by assessing GFP expression and *Gadd45*γ mRNA levels (Figure S1A-C). Neither group showed anatomical or histological brain abnormalities. Two weeks after stereotaxic surgery, before assessing cognitive function, we performed an open field test (Figure 1C) to verify whether Gadd45γ overexpression affects locomotor activity or anxiety-like behavior. Total distance travelled and the percentage of the time spent in the central zone were similar between groups (Figure 1D-F). Next, we assessed long-term memory in the object-place recognition test and contextual fear conditioning. Increasing Gadd45γ expression in the dHPC of young mice impaired preference for the displaced object 24h after learning (Figure 1G). This impairment was not due to altered habituation patterns during the training trial sessions or altered object exploratory behavior (Figure S1D,E). In contrast, Gadd45γOE mice showed intact long-term memory in contextual fear conditioning (Figure 1H). Both groups presented similar responses to shock administration (Figure S1F).

### 3.3. Gadd45γ overexpression disrupts activity-dependent CREB activation and gene expression

Considering that Gadd45γ regulates CREB activity (Brito et al., 2020), we next investigated whether Gadd45γ overexpression would impact CREB regulation. First, we assessed whether Gadd45γ overexpression in the hippocampus of young adult mice (Figure 2A) affects the levels of phosphorylated CREB in baseline conditions. In agreement with a role for Gadd45γ in signaling pathways that regulate CREB activation (Brito et al., 2020), overexpression of Gadd45γ resulted in increased levels of phosphorylated CREB (Figure 2B,C). Next using primary hippocampal cultures (Figure S1G,H) we analyzed the phosphorylation of CREB in baseline conditions and in response to increased neuronal activity (Figure 2D). Similar to our *in vivo* findings (Figure 2A-C), Gadd45γ overexpression in primary hippocampal cultures led to increased levels of phosphorylated CREB in basal conditions. Moreover, this effect appeared to blunt a response to neuronal activity; CREB phosphorylation in response to neuronal activity did not reach controls levels (Figure 2E,F). We next assessed the expression of the CREB-dependent genes *Arc, FosB, c-Fos, Egr1 and Npas4* (Impey et al., 2004; Rao-Ruiz et al., 2019) in basal conditions and upon neuronal activity (Figure 2G). Hippocampal neuronal cultures infected with rAAV-Gadd45γOE revealed disrupted CREB-dependent gene expression in response to increased neuronal activity compared to control conditions (Figure 2H-L). This set of experiments shows that increasing Gadd45γ above physiological levels in hippocampal neurons disrupts CREB phosphorylation and gene expression required for memory formation. Taken together, these findings demonstrate that an increase in hippocampal Gadd45γ levels disrupts CREB regulation, the expression of memory-related genes and cognitive function.

**Figure 2.**
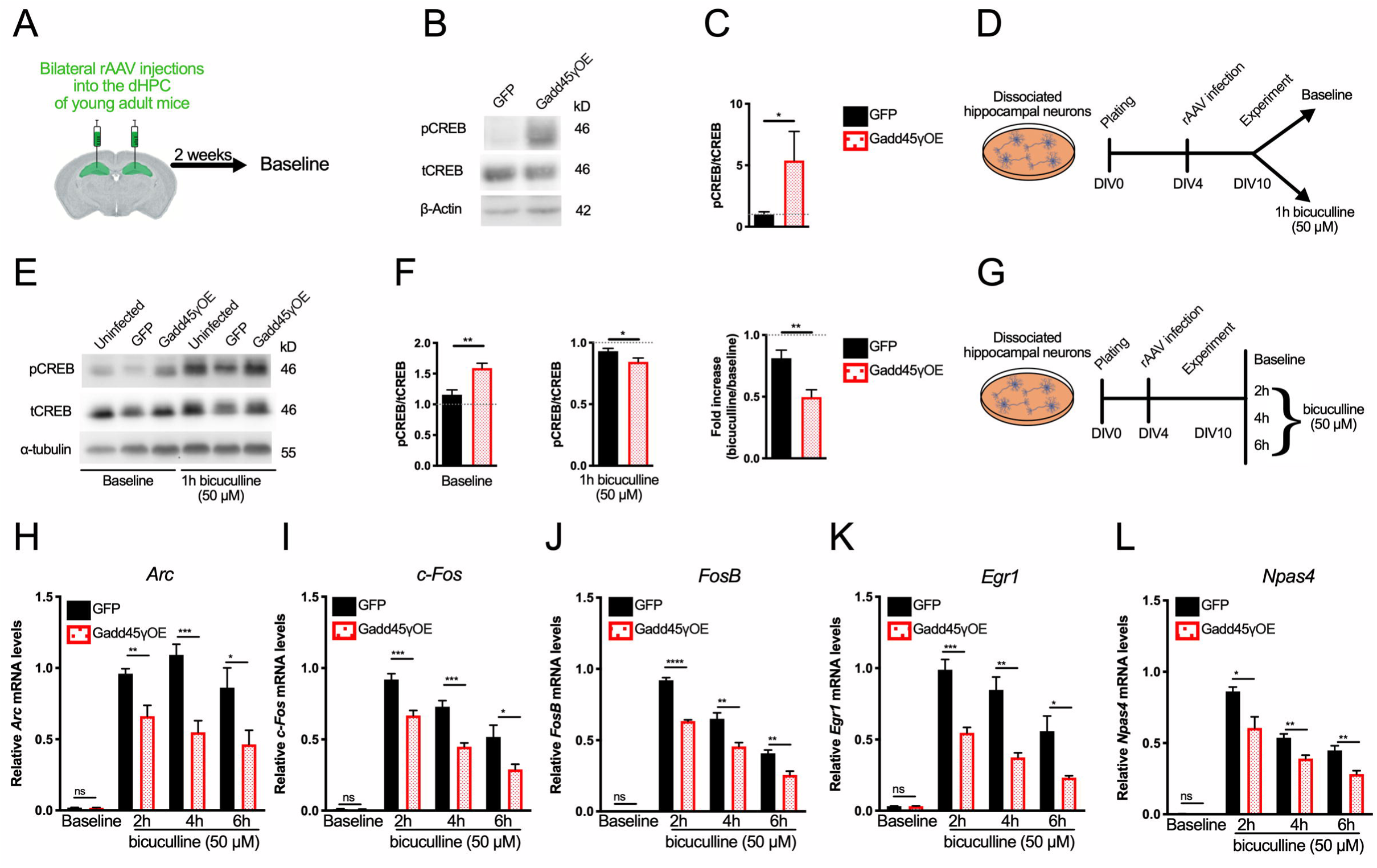
Increased Gadd45γ expression dysregulates hippocampal CREB phosphorylation and activity-dependent gene expression. **A)** Schematic representation of the experimental design for western blot analysis of CREB activation *in vivo*. **B)** Representative immunoblot scans of hippocampal tissue from young adult mice infected with rAAVs expressing GFP or Gadd45γOE using phosphospecific (pCREB) and total CREB (tCREB) antibodies. **C)** Immunoblot quantification shown as ratios of phosphorylated/total protein normalized to GFP control (N=5-6). **D)** Schematic representation of the experimental design for western blot analysis of CREB activation in mouse hippocampal cultures. **E)** Representative immunoblot scans of hippocampal cultures infected with rAAV expressing GFP or Gadd45γOE using phosphospecific and total CREB antibodies. **F)** Immunoblot quantification shown as ratios of phosphorylated/total protein normalized to uninfected control in baseline or bicuculline-treated conditions (left and middle graphs) or pCREB fold increase (bicuculline/baseline) for each condition and normalized to uninfected control (right graph) (N=7-8 independent cell preparations). **G)** Schematic representation of the experimental design for qRT-PCR analysis of the expression of CREB-dependent genes (N=5-6 independent cell preparations) **H)** *Arc*, **I)** *c-Fos* **J)** *FosB* **K)** *Egr1* and **L)** *Npas4* in hippocampal cultures. Hippocampal cultures were harvested at baseline conditions or after 2h, 4h, or 6h of bicuculline treatment. *p<0.05, **p<0.01, ***p<0.001 and ****p<0.0001 by two-tailed Student’s t-test. ns: not significant. Error bars represent SEM.

## 4. Discussion

This study suggested that human aging is associated with increased hippocampal Gadd45γ expression. Together with our previous findings (Brito et al., 2020) we showed that bidirectional dysregulation of hippocampal Gadd45γ levels in young adult mice negatively impacts cognitive function, CREB regulation and the expression of memory-related genes. Thus, implicating a requirement for tight control of Gadd45γ levels in brain function.

We observed that mimicking the human aging-related increase in Gadd45γ expression in the mouse hippocampus or in dissociated hippocampal neurons, promoted memory deficits and impairments in CREB-dependent gene transcription, respectively. Both the reduction or chronic enhancement of CREB function is known to lead to spatial memory deficits (Li et al., 2015; Pittenger et al., 2002; Viosca et al., 2009), as described in studies using CREB-deficient mutants (Pittenger et al., 2002) or mouse models that express constitutively active forms of CREB such as VP16-CREB (Viosca et al., 2009). Moreover, constitutive CREB activation has been identified as a possible contributing mechanism involved in Alzheimer’s disease (Muller et al., 2011). In our study Gadd45γ overexpression induced increased levels of phosphorylated CREB in basal conditions both in the mouse hippocampus and in hippocampal cultures. Although this remains to be investigated, the observed increase in Gadd45γ expression levels in the aged human hippocampus, may also result in chronic elevations in the activated form of CREB. Our experiments in hippocampal cultures indicate that increased levels of phosphorylated CREB already in basal conditions impair a response to neuronal activity and result in deficits in the expression of plasticity-related genes. Thus, providing a possible mechanism for impaired cognitive function in conditions of elevated Gadd45γ expression like in our model and possibly in the hippocampus of aged individuals. Together with our previous findings, which showed that Gadd45γ knockdown leads to impairments in CREB activation and associated gene expression (Brito et al., 2020), this data suggests that proper cellular function requires the tight regulation of Gadd45γ levels. These findings are in agreement with another study showing that either Gadd45γ loss- or gain-of-function disrupts neural development (Kaufmann and Niehrs, 2011).

The deficits in memory were task-specific; young adult mice expressing Gadd45γ above physiological levels presented selective long-term memory impairments in object place-recognition memory but not in contextual fear conditioning. Intriguingly, similar results have been found in aged mice and humans. It has been described that aged mice (Kennard and Woodruff-Pak, 2011) and humans (Battaglia et al., 2018; Foster et al., 2012; Leal and Yassa, 2015) are more likely to display deficits in forms of recognition memory than in contextual fear conditioning. The reasons for the selective impairment may be attributed to the characteristics of the tasks; despite hippocampal dysfunction in response to aging or Gadd45γ overexpression, mice may still be able to form and store the association between a highly salient stimulus (novel context) and a foot-shock. Similar findings have been described for other models of impaired hippocampal function. Namely, in a mouse model of Rett syndrome (Gulmez Karaca et al., 2018) and Alzheimer’s disease (Corcoran et al., 2002). In the later, contextual fear conditioning impairments were only present when the salience of the context was reduced.

Aberrant gene transcription patterns occur as a consequence of aging in the hippocampus (Burger, 2010; Ianov et al., 2017; Verbitsky et al., 2004). These changes do not overly correlate across species (Bishop et al., 2010; Loerch et al., 2008; Zahn et al., 2007), thus limiting the translational potential of animal models. Studies comparing cross-species alterations in gene expression generally focus on shared changes. The similar consequences of bidirectional dysregulation of Gadd45γ expression levels suggest that this approach may neglect functionally relevant and seemingly disparate age-associated transcription changes. Using *in vivo* and *in vitro* models we show that hippocampal levels of Gadd45γ are tightly regulated and that either a decrease (Brito et al., 2020) or an increase in Gadd45γ can dysregulate plasticity-associated gene expression and cause cognitive impairments. Notably, previous studies that performed transcriptomic analysis of human hippocampal tissue throughout adulthood also suggested an age-associated increase in Gadd45γ expression (Kang et al., 2011; Pletikos et al., 2014). Accordingly, our findings illustrate a scenario in which diverging age-related transcriptional programs in mice and humans result in converging phenotypes.

Taken together, our results demonstrate the requirement for tight control of Gadd45γ levels in memory formation and further implicate Gadd45γ as a molecular candidate that may underlie cognitive impairments in aging-associated pathological conditions.

## Acknowledgements

We thank I. Bu□nzli-Ehret for the preparation of primary hippocampal cultures and Stephanie Zeuch for comments to the manuscript. This work was supported by the Deutsche Forschungsgemeinschaft (DFG) [SFB 1134 (C01), OL 437/1 to A.M.M.O.], by Chica and Heinz Schaller foundation [fellowship to A.M.M.O.] and by Santa Casa da Misericórdia Mantero Belard Award [MB-07-2018 to L.V.L.].

