## Supplemental Information for "Modeling human age-associated increase in Gadd45γ expression leads to spatial recognition memory impairments in young adult mice"

by David VC Brito *et al*.

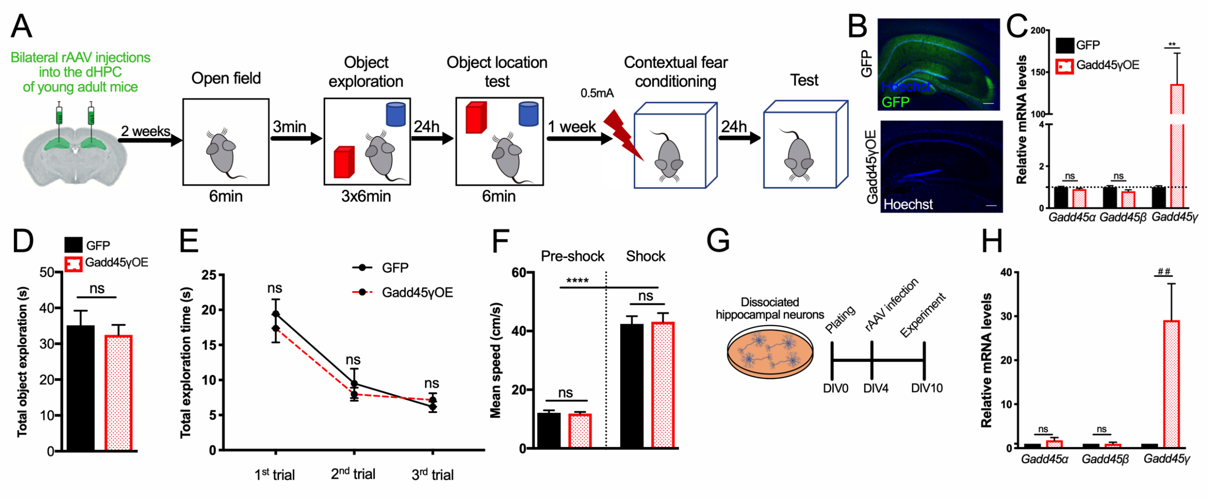


**Supplementary Figure 1.** *In vivo* and *in vitro* characterization of Gadd45γ overexpression. **A)** Schematic representation of the experimental design used for the behavioral analysis. **B)** Representative images of the dorsal hippocampus injected with viruses leading to either Gadd45γ-specific overexpression (Gadd45γOE) or the control expression of GFP, 4 weeks after stereotaxic surgery. Scale bar=100µm. **C)** qRT-PCR analysis of *Gadd45α, Gadd45β* and *Gadd45γ* expression levels in dHPC tissue infected with GFP or Gadd45γOE (N=6). A two-tailed unpaired Student's t-test was used. **D)** Total object exploration time during the training session of the object-location task. One-way repeated measure ANOVA was used (N=9). **E)** Total object exploration time during each trial of the training session compared to the first trial. Similar habituation patterns were observed between groups (N=9). Two-tailed unpaired Student's t-test was used. **F)** Mean speed during the different phases of the contextual fear conditioning training, showing similar performance between groups. A one-way ANOVA followed by a Bonferroni's Multiple Comparisons Test was used (N=9). **G)** Schematic representation of the experimental design used for gene expression analysis. **H)** qRT-PCR analysis of *Gadd45α*, *Gadd45β*, and *Gadd45γ* expression levels in cultured hippocampal cells infected with GFP or Gadd45γOE in baseline conditions (N=6 independent cell preparations). Data are normalized to the uninfected control. Kruskal-Wallis Test followed by a Dunn's Multiple Comparisons Test was used. $^{\#\#}p<0.01$, $^{**}p<0.01$ and $^{****}p<0.0001$. ns: not significant. Error bars represent SEM.
